# The first *de novo* HiFi genome assemblies for three clownfish-hosting sea anemone species (Anthozoa: Actiniaria)

**DOI:** 10.1101/2024.11.13.623406

**Authors:** Aurélien De Jode, Benjamin M. Titus

**Affiliations:** Department of Biological Sciences, University of Alabama, Tuscaloosa, AL, USA; Dauphin Island Sea Lab, Dauphin Island, AL, USA

**Keywords:** Cnidaria, Actinioidea, Symbiosis

## Abstract

The symbiosis between clownfish and giant tropical sea anemones (Order Actiniaria) is one of the most iconic on the planet. Distributed on tropical reefs, 28 species of clownfishes form obligate mutualistic relationships with 10 nominal species of venomous sea anemones. Our understanding of the symbiosis is limited by the fact that most research has been focused on the clownfishes. Chromosome scale reference genomes are available for all clownfish species, yet there are no published reference genomes for the host sea anemones. Recent studies have shown that the clownfish-hosting sea anemones belong to three distinct clades of sea anemones that have evolved symbiosis with clownfishes independently. Here we present the first high quality long read assemblies for three species of clownfish hosting sea anemones belonging to each of these clades: *Entacmaea quadricolor, Stichodactyla haddoni, Radianthus doreensis*. PacBio HiFi sequencing yielded 1,597,562, 3,101,773, and 1,918,148 million reads for *E. quadricolor, S. haddoni*, and *R. doreensis*, respectively. All three assemblies were highly contiguous and complete with N50 values above 4Mb and BUSCO completeness above 95% on the Metazoa dataset. Genome structural annotation with BRAKER3 predicted 20,454, 18,948 and 17,056 protein coding genes in *E. quadricolor, S. haddoni* and *R. doreeensis* genome, respectively. These new resources will form the basis of comparative genomic analyses that will allow us to deepen our understanding of this mutualism from the host perspective.

**Significance:** Chromosome-scale genomes are available for all 28 clownfish species yet there are no high-quality reference genomes published for the clownfish-hosting sea anemones. The lack of genomic resources impedes our ability to understand evolution of this iconic symbiosis from the host perspective. The clownfish-hosting sea anemones belong to three clades of sea anemones that have evolved mutualism with clownfish independently. Here we assembled the first high-quality long-read genomes for three species of host sea anemones each belonging to a different host clade: *Entacmaea quadricolor, Stichodactyla haddoni, Radianthus doreensis*. These resources will enable in depth comparative genomics of clownfish-hosting sea anemones providing a critical perspective for understanding how the symbiosis has evolved. Finally, these reference genomes present a significant increase in the number of high-quality long-read genome assemblies for sea anemones (11 currently published) and double the number of high-quality reference genomes for the sea anemone superfamily Actinoidea.

## Introduction

Among all mutualistic symbioses none are more recognizable than the association between clownfishes (also called anemonefishes: Subfamily Amphiprioninae) and their giant host sea anemones (Anthozoa: Actiniaria). Distributed broadly on tropical coral reefs throughout the Indian and Pacific Oceans, 28 species of clownfishes have rapidly radiated to form obligate mutualistic relationships with 10 nominal species of venomous sea anemones (Litsios et al., 2012; Gaboriou et al., 2024; Titus et al. 2024). Famously, clownfishes have evolved to live unharmed in their toxic and venomous hosts benefiting from their association with sea anemones by receiving shelter against predators (Fautin 1991), protection for their externally brooded eggs (Fautin & Allen 1992), removal of external parasites (Lubbock 1981) and potentially even nourishment (Verde et al., 2015). In turn, sea anemones are granted reciprocal protection against predation (Dunn 1981, Godwin & Fautin, 1992, Holbrook & Schmitt, 2005), novel sources of nitrogen and carbon from fish excrement (Roopin et al. 2008, Roopin & Chadwick, 2009, Cleveland et al., 2011), and increased oxygenation and gas transfer as clownfishes move through their tentacles (Szczebak et al. 2013).

The symbiosis has attracted a great deal of popular and scientific attention. The small size of clownfishes and their well-defined hierarchical social groups, reproductive biology, and amenability to aquaculture have made them tractable systems for understanding fundamental biological processes. They are now considered an emerging model organism for a wide range of research (Roux et al., 2020; Laudet & Ravasi, 2022). Chromosome-level reference genomes are now available for all 28 clownfish species (Marcionetti et al., 2019; Marcionetti & Salamin, 2023; Gaboriau et al., 2024) and comparative analyses are revealing the first candidate genes likely to be associated with the ability of clownfishes to remain protected from toxic sea anemone hosts (Marcionetti et al., 2019; Marcionetti & Salamin, 2023).

Our evolutionary understanding of the symbiosis, however, is limited by the fact that most research and genomic resources have been focused on clownfishes. No reference genomes exist for any of the 10 clownfish-hosting sea anemones. This perspective is expected to be especially critical as recent work has demonstrated that the host sea anemones are driving phenotypic convergence in clownfishes (Gaboriau et al., 2024), and that host sea anemones have diversification times that are broadly concomitant with the clownfish radiation (De Jode et al. 2024). These findings highlight the importance of the host sea anemones for a holistic understanding of the entire symbiosis. High-quality reference genomes for the host sea anemones thus hold the potential to identify genomic regions associated with establishing and maintaining mutualism with clownfishes, and other genomic consequences of a symbiotic lifestyle.

Interestingly, unlike the 28 described species of clownfishes which form a monophyletic clade, the 10 clownfish hosting sea anemones belong to three distinct lineages within the sea anemone superfamily Actinioidea that have evolved symbiosis with clownfishes (Titus et al. 2019; 2024; De Jode et al. 2024). These have been coined *Entacmaea*, Stichodactylina, and Heteractina. The clade *Entacmaea* contains only the bubble-tip sea anemone, *Entacmaea quadricolor* (Fig. 1A), which appears to be a species complex (Titus et al. 2019; De Jode et al. 2024). Clade Stichodactylina contains five clownfish hosting sea anemones species: *Cryptodendrum adhaesivum, Radianthus magnifica, Stichodactyla gigantea, S. haddoni* (Fig. 1B), and *S. mertensii*, as well as other non-host species (e.g. De Jode et al. 2024). Clade Heteractina includes *Heteractis aurora, Radianthus crispa, R. doreensis* (Fig. 1C), and *R. malu* (Titus et al. 2019; De Jode et al. 2024). The three independent clades of host sea anemones provide a natural starting point for sequencing and assembling high-quality reference genomes.

**Figure 1.**
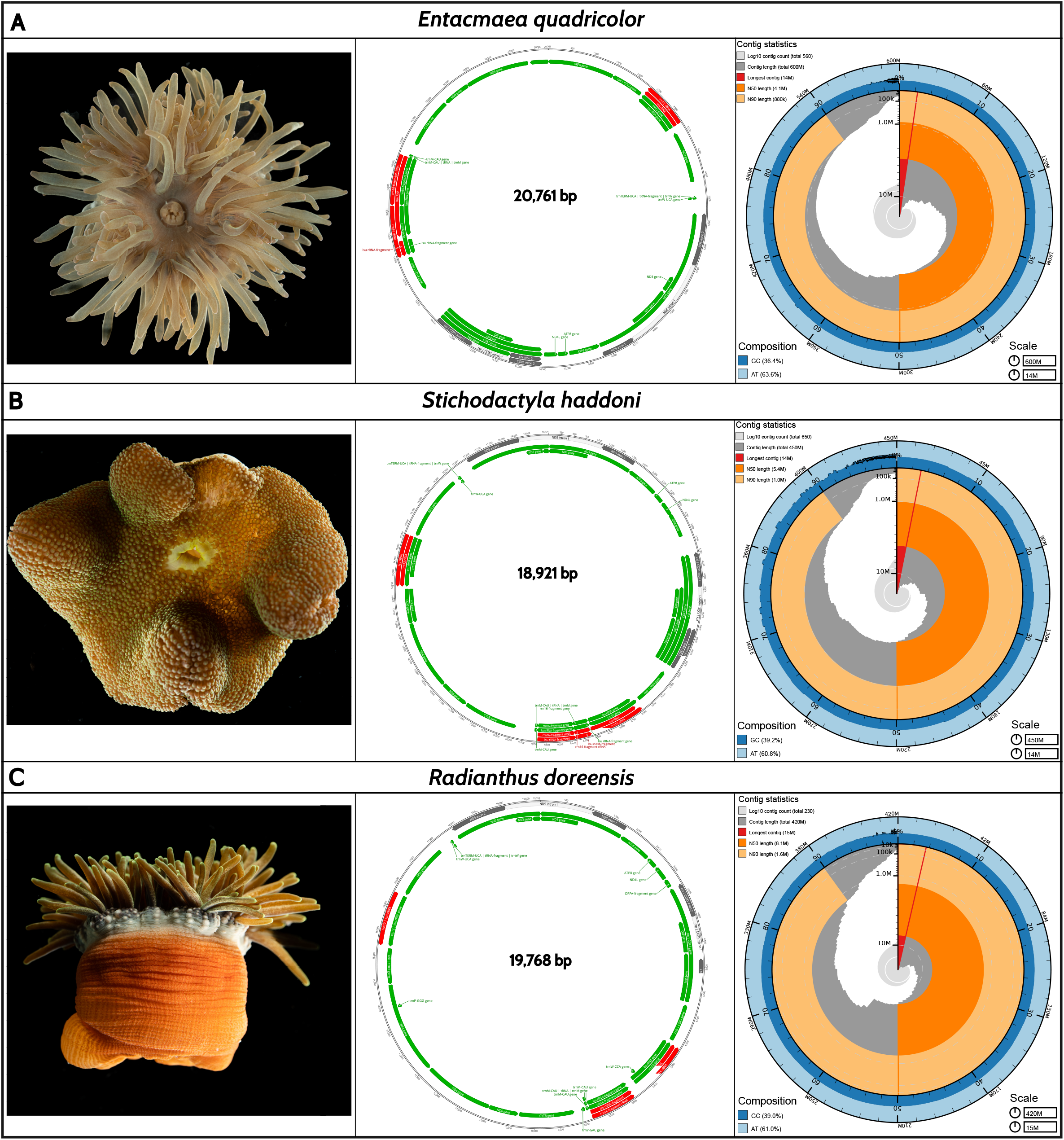
Long-read mitochondrial and nuclear genome assemblies for the clownfish hosting sea anemone species A) *Entacmaea quadricolor*, B) *Stichodactyla haddoni*, and C) *Radianthus doreensis*. Pictures of each species are the individuals sequenced in this study. Circularized and annotated mitogenomes were visualized with Geneious version 2024.0. BlobToolKit snailplots representing nuclear genome assembly statistics including genome size, largest scaffold length, N50 and N90 lengths, and GC composition. Snailplots are divided into 1,000 size-ordered bins around the circumference with each bin representing 0.1% of the assembly length. The distribution of contig lengths is shown in dark gray with the plot radius scaled to the longest contig present in the assembly (the smallest of the three arcs, also shown in red). Next two sectors in size (orange and pale-orange arcs) show the N50 and N90 contig lengths, respectively. The pale gray spiral shows the cumulative scaffold count on a log scale with white scale lines showing successive orders of magnitude. The dark-blue and pale-blue area around the outside of the plot shows the distribution of GC, AT, and N percentages in the same bins as the inner plot.

Here we present the first high quality long reads assemblies for three species of clownfish hosting sea anemones: *Entacmaea quadricolor, Stichodactyla haddoni*, and *Radianthus doreensis* (Fig. 1). These genomes will form the basis for in depth comparative genomics between clownfish hosting and non-hosting sea anemones and deepen our understanding of this iconic symbiosis.

## Results and Discussion

A total of 1,597,562 (∼ 20 Gbp), 3,101,773 (∼ 39 Gbp) and 1,918,148 (∼26 Gbp) HiFi reads were sequenced for *E. quadricolor, S. haddoni* and *R. doreensis*, respectively. For each species, mitochondrial genomes were assembled, circularized, and annotated. The total lengths of the newly assembled mitochondrial genomes were 20,761 bp, 18,921 bp, and 19,768 bp for *E. quadricolor, S. haddoni and R. doreensis* (Fig. 1), respectively. These are similar to the size of the mitogenomes used as references to identify mitochondrial reads. A characteristic feature of Hexacorallian mitogenomes is the presence of a self-splicing intron in the NAD5 gene (Feng et al., 2023). Interestingly, MitoFinder (the default annotation tool of MitoHiFi) did not retrieve that feature in our mitogenomes, but we were able identify and retrieve the NAD5 intron using MITOS in all species. Feng et al. (2023) identified a conserved mitochondrial gene order in the Order Actiniaria. Using the annotation conducted in GeSeq we confirmed that our three mitogenomes shared that same order (Act1GO sensu Feng et al. 2023).

Nuclear genome size estimates from GenomeScope were comparable between *S. haddoni* (339 Mbp) and *R. doreensis* (330 Mbp). The assembled genomes of *S. haddoni* and *R. doreensis* spanned 447 Mb and 418 Mb, respectively (supplementary table S1), both exceeding GenomeScope estimates by approximately 100 Mb. *Entacmaea quadricolor* had the largest estimated genome size at 578 Mbp, which was close to our final total assembly length of 599 Mb (supplementary table S1). Our genome size estimate for *E. quadricolor* was smaller than the 863 Mbp estimated using flow cytometry (Adachi et al., 2017), but higher than the 428 Mb assembly available on GenBank (GCA_024752375.1). The discrepancy between our PacBio assembly and the publicly available assembly is likely due to the latter genome being assembled using Illumina short reads.

Heterozygosity rates were comparable between *R. doreensis* (het: 0.767%) and *S. haddoni* (het: 0.855%) but were higher in *E. quadricolor* (het: 1.49 %). Merqury k-mer spectra plots revealed the presence of haplotypic duplications in each primary assembly (supplementary fig. 1). For each species, the assembly we obtained after using the purge_dups algorithm with manual cutoffs and contigs identified as repeats added back into the assembly, showed the best compromise between under and over purging and was kept for further analysis (supplementary fig. 1). A total of 17, 8 and 8 contigs were identified as contaminants and removed from the *E. quadricolor, S. haddoni* and *M. doreensis* assemblies, respectively (supplementary fig. 2). One mitochondrial contig was also identified and removed from each assembly.

Final genome assemblies were highly contiguous and comprised of 558, 648 and 230 contigs for *E. quadricolor, S. haddoni* and *R. doreensis*, respectively. N50 values were roughly 4 Mb (*E. quadricolor*), 5.3 Mb (*S. haddoni*), and 8 Mb (*R. doreensis*) (supplementary table 1). BUSCO completeness scores for the Metazoa dataset were 95.6 % for *E. quadricolor* and *S. haddoni* and 95.4 % for *R. doreensis* (supplementary table 1). A total of 20,454, 18,948 and 17,056 protein coding genes were predicted by the BRAKER3 annotation pipeline for *E. quadricolor, S. haddoni* and *R. doreensis* respectively (Fig. 2). For each species the total number of predicted proteins were comparable to the number of predicted proteins for other members of Order Actiniaria (Fig. 2).

**Figure 2.**
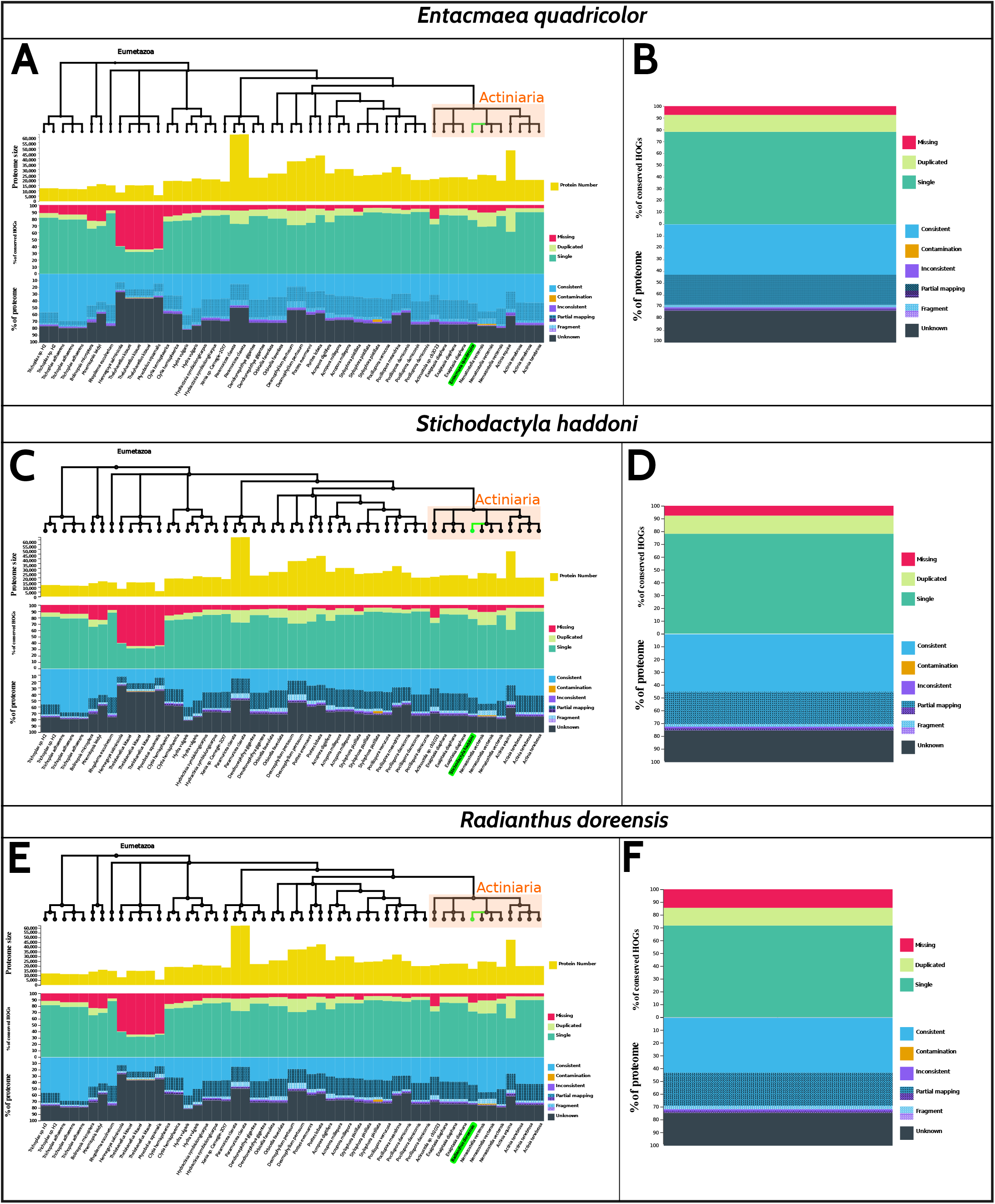
OMArk quality assessment of genome annotations. A, C, &E) OMArk comparison plots showing genome annotation statistics for *Entacmaea quadricolor, Stichodactyla. haddoni* and *Radianthus doreensis* in reference to other members of Eumetazoa. Positions of the clownfish-hosting sea anemones sequenced in this study are highlighted in green. List of available proteome accessions used for the comparisons in Supplementary material. B, D, & F) detailed proteome assessment for each species of clownfish-hosting sea anemones sequenced in this study.

The OMArk completeness percentages, based on Hierarchical Orthologous Groups (HOGs) of genes in Eumetazoa, were 92.6%, 92.35 % and 85.53 % for *E. quadricolor, S. haddoni* and *R. doreensis*, respectively (Fig. 2, supplementary text). The OMArk consistency assessment was very similar among our three species and other members of Order Actiniaria (Fig. 2, supplementary text). OMArk consistency (the gene family is known to exist in the Eumetazoa lineage) scores 70.16 %, 72.27 % and 71. 56 % for *E. quadricolor, S. haddoni* and *R. doreensis*, respectively. No protein sequence was identified as a contaminant in any of our species. The lower amount of protein coding genes predicted in *S. haddoni* and *R. doreensis* and the lower level of completeness in *R. doreensis* are likely due to the lower amount of RNA seq data available for those species (supplementary text). The ratio of mono- to multi-exonic transcripts were 0.34, 0.28 and 0.34 for *E. quadricolor, S. haddoni* and *R. doreensis* respectively.

## Conclusion

Prior to this study, no genomes had been published for the clownfish-hosting sea anemones, although chromosome level genomes are available for all 28 species of clownfishes (Gaboriau et al. 2024). The three high-quality genomes we’ve assembled here using PacBio HiFi reads belong to species from the three clades that have independently evolved mutualism with clownfishes (Titus et al., 2019; De Jode et al., 2024). These will thus become critical new resources for comparative genomics and our understanding of clownfish-hosting sea anemones diversification and of the evolution of the iconic mutualism. Finally, only 11 long-read genomes have been assembled and published for Order Actiniaria. Our genomes represent a significant increase in the number of genomic resources available for understanding sea anemone biology and evolution broadly.

## Material and Methods

Live sea anemones were purchased via the ornamental aquarium trade through Quality Marine Inc. to ensure the geographic origin of each animal. We acquired three species: the bubble tip sea anemone *Entacmaea quadricolor* (Fiji), Haddon’s carpet sea anemone *Stichodactyla haddoni* (Vietnam), and the long-tentacled sea anemone *Radianthus doreensis* (Vietnam). Fresh or flash frozen pedal disc tissue was thinly sliced from each anemone to avoid endosymbiotic dinoflagellate contamination and used for DNA extractions. High Molecular Weight (HMW) DNA was extracted using PacBio Nanobind® tissue kits following manufacturer instructions with the exception that the quantity of proteinase K was increased to 40μL. DNA samples were purified with Monarch Nucleic Acid Purification Kit (New England BioLabs, Ipswich, Massachusetts) prior to shearing on a Megaruptor 2 (Diagenode, Denville, NJ), at 20kb setting.

Libraries were constructed and sequenced at Maryland Genomics at the Institute for Genome Sciences, University of Maryland School of Medicine. Samples were prepared using SMRTbell Prep Kit 3.0 (Pacific Biosciences, Menlo Park, CA) with barcoded adaptors, and subsequently size-selected on BluePippin instrument (Sage Science, Beverly, MA) to remove DNA fragments less than 10kb in size, followed by an equimolar pooling. The pooled libraries were bound to Revio polymerase, then sequenced with a Revio Sequencing Plate and flow cell on a PacBio Revio platform (Pacific Biosciences, Menlo Park, CA). Mitochondrial genomes were assembled from raw HiFi reads using the MitoHiFi v2. (Uliano-Silva et al., 2023) pipeline with default parameters except for the –max-read-len option that was set to 2. Mitochondrial reads were identified for *E. quadricolor, S. haddoni*, and *R. doreensis* using the mitogenomes of *E. quadricolor* (Genbank NC_049066.1); *Heteractis aurora* (Genbank NC_047219.1) and *S. haddoni* (MW760873.1, Johansen et al., 2021), respectively. Mitochondrial genome annotation was performed using both MitoFinder (Allio et al., 2020) and MITOS (Bernt et al., 2013). A final annotation was conducted using GeSeq (Tillich et al., 2017) (see explanations in Results & Discussion section). Mitogenomes were visualized using Geneious version 2024.0 (https://www.geneious.com) created by Biomatters.

Genome sizes and heterozygosities were estimated using GenomeScope 2.0 (Ranallo-Benavidez et al., 2020) based of k-mer histogram frequencies generated by Jellyfish v2.3.0 (Marçais & Kingsford, 2011) for a k-mer size of 21bp (Vulture et al. 2017). Nuclear genomes were assembled using Hifiasm v0.19.8-r603 (Cheng et al., 2021) and the first set of partially phased contigs was used for the following analysis.

Haplotypic duplications, were removed from the assemblies using the purge_dups v1.2.5 (Guan et al., 2020) algorithm with the following customs cutoffs: (3, 18, 18, 19, 19, 180), (20, 54, 54, 55, 55, 240) and (7, 39, 39, 40, 40, 200) for *E. quadricolor, S. haddoni* and *M. doreensis*, respectively. Contigs classified as repeats by purge_dups were added back to the assemblies. The Blobtoolkit (Laetsch & Blaxter, 2017, Challis et al. 2020) v3.4.3 pipeline was then used to assess the presence of contaminant contigs in each assembly. Mitochondrial contigs were removed from decontaminated nuclear contigs using MitoHiFi v2. (Uliano-Silva et al., 2023) with the mitochondrial genome assembled from the reads as a reference.

Genomes were annotated using the BRAKER3 pipeline v.3.0.8 (Gabriel et al. 2024) specifying a BUSCO lineage (--busco_lineage=metazoa_odb10). The pipeline uses several sub-pipelines and third-party tools (Stanke et al., 2006, Gotoh, 2008, Stanke et al., 2008, Iwata & Gotoh, 2012, Simão et al., 2015, Buchfink et al., 2015, Hoff et al., 2016, Hoff et al., 2019, Kovaka et al., 2019, Pertea & Pertea, 2020, Brůna et al., 2021, Gabriel et al., 2021, Huang & Li, 2023, Li, 2023, Brůna et al., 2024) to predict highly reliable genes in reference genome using protein and RNA-seq data. RNA-seq data (see SI for list of accession numbers) were downloaded from Genbank using the fastq-dump function of the SRA-toolkit v.3.0.7 (SRA Toolkit Development Team) and reads were mapped to the genome using HISAT2 v.2.2.1 (Kim et al. 2019) with the --dta option. The Metazoa partition of the OrthoDB v11 protein database was used as evidence for BRAKER 3 (Kuznetsov et al., 2023). To compute the ratio of mono- to multi-exon genes we used the analyze_exons.py script from the GALBA github repository (https://github.com/Gaius-Augustus/GALBA). Annotation quality was assessed with use OMArk v.0.3.0 (Nevers et al., 2022) on the OMArk public webserver (https://omark.omabrowser.org/home/). Genome quality was assessed for the final assembly and at several intermediate steps during the assembly process. Contiguity and completeness metrics were computed using gfastats v1.3.6 (Formenti et al., 2022) and BUSCO v5.6.1 (Manni et al. 2021). Completeness, quality and levels of haplotypic duplication were assessed using Merqury (Rhie et al. 2020) based on HiFi reads.

## Acknowledgments

We thank members of the Titus Lab for help with sample processing and Rebecca Varney for advice on high molecular weight DNA extractions. We thank Maryland Genomics at the Institute for Genome Sciences, University of Maryland School of Medicine for library preparation and sequencing. We also thank Giulio Formenti and Jonathan M. Wood for helpful discussion regarding the purging of haplotypic duplications. Funding for this work was provided by the University of Alabama and National Science Foundation Grant DEB-1934274 to BMT.

## Data Availability

The HiFi PacBio sequencing reads, are deposited in NCBI under the BioProject accession numbers PRJNA1076568. The final genome assembly and genome annotation will be deposited under the same BioProject upon publication.

## Notes

### Competing Interest Statement

The authors have declared no competing interest.

### Summary of Updates

Better pdf formatting so that legends are placed above their figure.

