## Supplementary Material for "The first *de novo* HiFi genome assemblies for three clownfish-hosting sea anemone species (Anthozoa: Actiniaria)"

RNA-seq data accession numbers……………………………………………………..Page 4
OMArk detailed results………………………………………………………………..Page 5 to 9
OMArk proteome accessions…………………………………………………………..Page 10

**Figure S1.** Merqury K-mer spectra plots. Merqury copy number spectrum plots for each genome after Hifiasm assembly (left plot), after one round of haplotigs removal with purge_dups and manual cuttoffs (middle plot) and after adding back the repeats (right plot). Each plot tracks the multiplicity of each k-mer found in the Hifi read set and colors it by the number of times it is found in a given assembly. For example, k-mers found only one time in the assembly are represented by the red line that forms 2 peaks of multiplicity. The first peak represents 1-copy (heterozygous) k-mers in the genome, and the second peak represents 2-copy k-mers originating from homozygous sequence or haplotype-specific duplications. Depth of sequencing coverage determines where these peaks appear. The assembly k-mers absent from the read set are plotted as a bar at zero multiplicity, colored by the copy numbers found in the assembly. Missing single copy k-mers (read-only) are plotted in black, and k-mers present more than one time in the assembly are represented in blue, green, purple and orange.

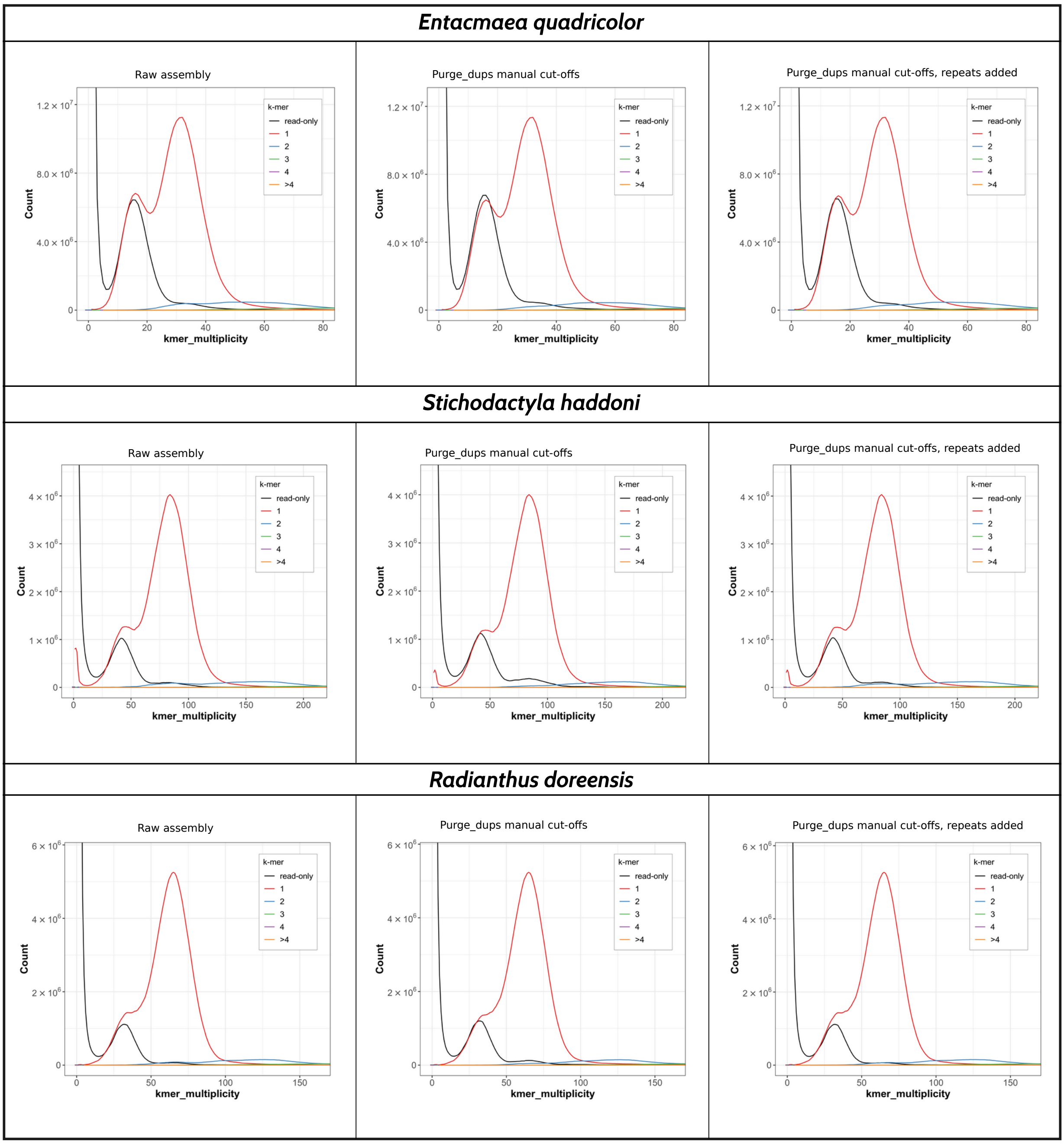

**Figure S2.** BlobToolKit GC-coverage plot of assemblies before contaminants removal. Black circle indicates contigs that were removed for being likely contaminants. Contigs are colored by phylum. Circles are sized in proportion to contig length. Histograms show the distribution of contig length sum along each axis. Black circles indicate the positions of the contigs that have been removed for being contaminants.

| Entacmaea quadricolor | Stichodactyla haddoni | Radianthus doreensis |
| --- | --- | --- |
| 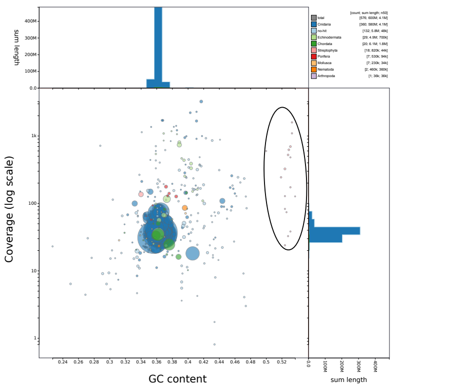  | 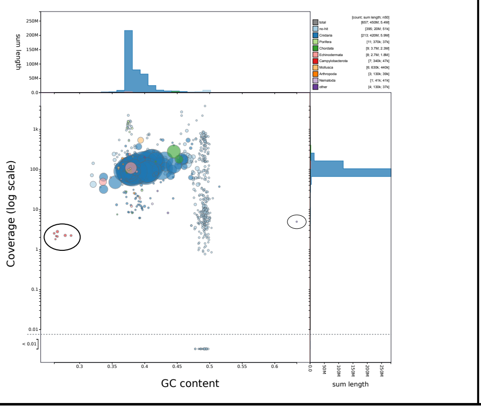  | 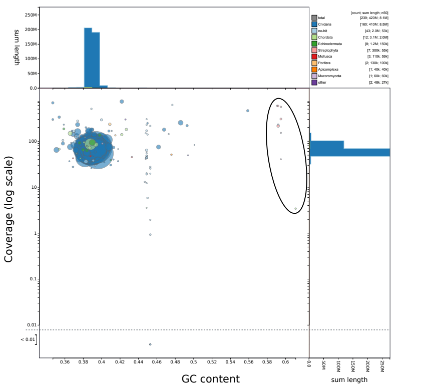  |

**List of RNA-seq data accession number used in the annotation pipeline :**

*Entacmaea quadricolor*: ERR2045166, ERR2045167, ERR2045168, ERR2045169, ERR2045170, ERR2045171, SRR14307520, SRR14307527, SRR14307530, SRR21497998, SRR21497999, SRR21498000, SRR21498002, SRR21 498013, SRR21498014, SRR23908431, SRR23908432, SRR23908433, SRR23908434, SRR23908435, SRR23908436, SRR23908437, SRR23908441, SRR23908450, SRR23908451, SRR23908452, SRR23908453, SRR27756961, SRR27756962, SRR27756963, SRR27756964, SRR27756965, SRR27756966, SRR27756967, SRR27756968, SRR27756969, SRR27756970, SRR27756971, SRR27756972, SRR27756973, SRR27756974, SRR27756975, SRR27756976, SRR27756977, SRR27756978, SRR27756979, SRR27756980, SRR27756981, SRR27756982, SRR27756983, SRR27756984

*Stichodactyla haddoni*: SRR29215550, SRR29215551, SRR29215552, SRR23908438, SRR23908439, SRR23908440, SRR23908442, SRR21498006, SRR21498007, SRR14307523, SRR5397293

*Radianthus doreensis*: SRR29774726, SRR14115222, SRR14115223, SRR14115224, SRR14115225

### OMArk detailed results

*-Entacmaea quadricolor*

COMPLETENESS ASSESSMENT
------------
#This benchmark gives an estimate of the completeness of the gene set based on the presence or not of conserved genes of the target lineage.
#Conserved genes are defined using Hierarchical Orthologous Groups (HOGs) defined at a certain taxonomic clade, which is a proxy for the ancestral gene repertoire of this clade. HOGs are considered conserved if they have at least one gene in >80% of the extant species.
#Because representatives of these groups are expected to be present in the target species repertoire, the proportion of missing HOGs proxies the proportion of missing genes in the total gene repertoire of the target proteome.
#Ancestral genes used for this benchmark were in single copy in the selected ancestral lineage, but no assumption is made regarding their propensity to duplicate - they are not universal single copy genes. This benchmark reports the proportion of those genes that are found in multiple copies in target proteomes, and whether it corresponds to a known duplication event in descendants of this gene family (Expected) or not (Unexpected).

The clade used was: Eumetazoa
Number of conserved HOGs: 3255

#Results on conserved HOGs:
Single: 2548 (78.28%)
Duplicated: 466 (14.32%)
Duplicated, Unexpected: 439 (13.49%)
Duplicated, Expected: 27 (0.83%)
Missing: 241 (7.40%)

CONSISTENCY ASSESSMENT
------------------------
#This benchmark gives the proportion of annotated protein-coding genes in the query proteome that likely correspond to an actual protein-coding gene by comparing to the known gene families of the selected ancestral lineage.

##High-level categories
#Genes in the "Consistent" category correspond to a gene family known to exist in the selected lineage. Genes in the "Inconsistent" or “Contaminants" categories correspond to known gene families from different lineages. Such genes are deemed contaminants if more genes than expected by chance correspond to the same species. They are deemed Inconsistent if they correspond to other species seemingly at random. Genes are classified in the “Unknown” category if they do not share enough similarity with known gene families: they may be orphan genes or erroneous protein sequences.

##Subcategories
#Partial hit proteins are those that share similarity with proteins in known gene families on only part of their sequence: they can indicate poorly defined gene models, structurally divergent genes, or erroneous annotation.
#Fragmented proteins are those whose length is smaller than the proteins from the gene families they share similarity with (<50% median length): they are likely fragmentend sequences or erroneous annotations.

Number of proteins in the whole proteome: 20454

#Consistent lineage placements
Total Consistent: 14350 (70.16%)
Consistent, partial hits: 5215 (25.50%)
Consistent, fragmented: 427 (2.09%)

#Inconsistent lineage placements
Total Inconsistent: 590 (2.88%)
Inconsistent, partial hits: 330 (1.61%)

Inconsistent, fragmented: 34 (0.17%)

#Contaminants
Total Contaminants: 0 (0.00%)
Contaminants, partial hits: 0 (0.00%)
Contaminants, fragmented: 0 (0.00%)

#Unknown
Total Unknown: 5514 (26.96%)

SPECIES COMPOSITION
-------------------
#This benchmark gives an estimate of the species composition of the dataset, according to HOGs placement. It reports the clades most consistent with the taxonomic distribution of gene families where coding-genes for the query proteomes were placed. The species to which most of the proteins in the query proteome are consistent with is called "Main species." The others are potential contaminants.
#This section also lists the numbers of proteins that can be associated to each of these clades, based on the taxonomic placement of the gene families they share similarity with.

##Detected species

#Main species
Clade: Nematostella vectensis
Number of associated query proteins: 14940 (73.04%)

### *-Radianthus doreensis*

COMPLETENESS ASSESSMENT
------------
#This benchmark gives an estimate of the completeness of the gene set based on the presence or not of conserved genes of the target lineage.
#Conserved genes are defined using Hierarchical Orthologous Groups (HOGs) defined at a certain taxonomic clade, which is a proxy for the ancestral gene repertoire of this clade. HOGs are considered conserved if they have at least one gene in >80% of the extant species.
#Because representatives of these groups are expected to be present in the target species repertoire, the proportion of missing HOGs proxies the proportion of missing genes in the total gene repertoire of the target proteome.
#Ancestral genes used for this benchmark were in single copy in the selected ancestral lineage, but no assumption is made regarding their propensity to duplicate - they are not universal single copy genes. This benchmark reports the proportion of those genes that are found in multiple copies in target proteomes, and whether it corresponds to a known duplication event in descendants of this gene family (Expected) or not (Unexpected).

The clade used was: Eumetazoa
Number of conserved HOGs: 3255

#Results on conserved HOGs:
Single: 2333 (71.67%)
Duplicated: 451 (13.86%)
Duplicated, Unexpected: 426 (13.09%)
Duplicated, Expected: 25 (0.77%)
Missing: 471 (14.47%)

CONSISTENCY ASSESSMENT
------------------------

#This benchmark gives the proportion of annotated protein-coding genes in the query proteome that likely correspond to an actual protein-coding gene by comparing to the known gene families of the selected ancestral lineage.

##High-level categories
#Genes in the "Consistent" category correspond to a gene family known to exist in the selected lineage. Genes in the "Inconsistent" or “Contaminants" categories correspond to known gene families from different lineages. Such genes are deemed contaminants if more genes than expected by chance correspond to the same species. They are deemed Inconsistent if they correspond to other species seemingly at random. Genes are classified in the “Unknown” category if they do not share enough similarity with known gene families: they may be orphan genes or erroneous protein sequences.

##Subcategories
#Partial hit proteins are those that share similarity with proteins in known gene families on only part of their sequence: they can indicate poorly defined gene models, structurally divergent genes, or erroneous annotation.
#Fragmented proteins are those whose length is smaller than the proteins from the gene families they share similarity with (<50% median length): they are likely fragmentend sequences or erroneous annotations.

Number of proteins in the whole proteome: 17055

#Consistent lineage placements
Total Consistent: 12205 (71.56%)
Consistent, partial hits: 4396 (25.78%)
Consistent, fragmented: 456 (2.67%)

#Inconsistent lineage placements
Total Inconsistent: 470 (2.76%)
Inconsistent, partial hits: 253 (1.48%)
Inconsistent, fragmented: 23 (0.13%)

#Contaminants
Total Contaminants: 0 (0.00%)
Contaminants, partial hits: 0 (0.00%)
Contaminants, fragmented: 0 (0.00%)

#Unknown
Total Unknown: 4380 (25.68%)

SPECIES COMPOSITION
-------------------
#This benchmark gives an estimate of the species composition of the dataset, according to HOGs placement. It reports the clades most consistent with the taxonomic distribution of gene families where coding-genes for the query proteomes were placed. The species to which most of the proteins in the query proteome are consistent with is called "Main species." The others are potential contaminants.
#This section also lists the numbers of proteins that can be associated to each of these clades, based on the taxonomic placement of the gene families they share similarity with.

##Detected species

#Main species
Clade: Nematostella vectensis
Number of associated query proteins: 12675 (74.32%)

### -*Stichodactyl haddoni*

COMPLETENESS ASSESSMENT
------------
#This benchmark gives an estimate of the completeness of the gene set based on the presence or not of conserved genes of the target lineage.

#Conserved genes are defined using Hierarchical Orthologous Groups (HOGs) defined at a certain taxonomic clade, which is a proxy for the ancestral gene repertoire of this clade. HOGs are considered conserved if they have at least one gene in >80% of the extant species.
#Because representatives of these groups are expected to be present in the target species repertoire, the proportion of missing HOGs proxies the proportion of missing genes in the total gene repertoire of the target proteome.
#Ancestral genes used for this benchmark were in single copy in the selected ancestral lineage, but no assumption is made regarding their propensity to duplicate - they are not universal single copy genes. This benchmark reports the proportion of those genes that are found in multiple copies in target proteomes, and whether it corresponds to a known duplication event in descendants of this gene family (Expected) or not (Unexpected).

The clade used was: Eumetazoa
Number of conserved HOGs: 3255

#Results on conserved HOGs:
Single: 2545 (78.19%)
Duplicated: 461 (14.16%)
Duplicated, Unexpected: 432 (13.27%)
Duplicated, Expected: 29 (0.89%)
Missing: 249 (7.65%)

CONSISTENCY ASSESSMENT
-------------------------
#This benchmark gives the proportion of annotated protein-coding genes in the query proteome that likely correspond to an actual protein-coding gene by comparing to the known gene families of the selected ancestral lineage.

##High-level categories
#Genes in the "Consistent" category correspond to a gene family known to exist in the selected lineage. Genes in the "Inconsistent" or “Contaminants" categories correspond to known gene families from different lineages. Such genes are deemed contaminants if more genes than expected by chance correspond to the same species. They are deemed Inconsistent if they correspond to other species seemingly at random. Genes are classified in the “Unknown” category if they do not share enough similarity with known gene families: they may be orphan genes or erroneous protein sequences.

##Subcategories
#Partial hit proteins are those that share similarity with proteins in known gene families on only part of their sequence: they can indicate poorly defined gene models, structurally divergent genes, or erroneous annotation.
#Fragmented proteins are those whose length is smaller than the proteins from the gene families they share similarity with (<50% median length): they are likely fragmentend sequences or erroneous annotations.

Number of proteins in the whole proteome: 18946

#Consistent lineage placements
Total Consistent: 13693 (72.27%)
Consistent, partial hits: 4759 (25.12%)
Consistent, fragmented: 419 (2.21%)

#Inconsistent lineage placements
Total Inconsistent: 541 (2.86%)
Inconsistent, partial hits: 283 (1.49%)
Inconsistent, fragmented: 30 (0.16%)

#Contaminants
Total Contaminants: 0 (0.00%)
Contaminants, partial hits: 0 (0.00%)
Contaminants, fragmented: 0 (0.00%)

#Unknown
Total Unknown: 4712 (24.87%)

SPECIES COMPOSITION
-------------------
#This benchmark gives an estimate of the species composition of the dataset, according to HOGs placement. It reports the clades most consistent with the taxonomic distribution of gene families where coding-genes for the query proteomes were placed. The species to which most of the proteins in the query proteome are consistent with is called "Main species." The others are potential contaminants.
#This section also lists the numbers of proteins that can be associated to each of these clades, based on the taxonomic placement of the gene families they share similarity with.

##Detected species

#Main species
Clade: Nematostella vectensis
Number of associated query proteins: 14234 (75.13%)

**OMArk proteome accessions**

Trichoplax sp. H2 [UP000253843]
Trichoplax sp. H2 [NCBI GCA_003344405.1]
Trichoplax adhaerens [Ensembl ASM15027v1]
Trichoplax adhaerens [UP000009022]
Trichoplax adhaerens [NCBI GCF_000150275.1]
Bolinopsis microptera [NCBI GCF_026151205.1]
Mnemiopsis leidyi [Ensemble MneLei_Aug2011]
Rhopilema esculnetum [NCBI GCF_013076305.1]
Henneguya salminicola [NCBI GCA_009887335.1]
Thelohanellus kitauei [Ensembl ASM82789v1]
Thelohanellus kitauei [UP000031668]
Thelohanellus kitauei [NCBI GCA_000827895.1]
Myxobolus squamalis [NCBI GCA_010108815.2]
Clytia hemisphaerica [UP000594262]
Clytia hemisphaerica [Ensemble Clytia_hemisphaerica_genome_assembly]
Hydra vulgaris [UP000694840]
Hydra vulgaris [Ensemble Hudra_105_v3]
Hydractinia symbiolongicarpus [NCBI GCF_029227915.1]
Hydractinia symbiolongicarpus [Ensembl HSymV2.1]
Xenia sp. Carnegie-2017 [NCBI GCF_021976095.1]
Paramuricea clavata [UP001152795]
Paramuricea clavata [NCBI GCA_902702795.2]
Dendronephthya gigantea [NCBI GCF_00432835.1]
Dendronephthya gigantea [Ensembl DenGig_1.0]
Orbicella faveolata [NCBI GCF_002042975.1]
Orbicella faveolata [Ensembe ofav_dov_v1]
Desmophyllum pertusum [UP001163046]
Desmophyllum pertusum [NCBI GCA_029204205.1]
Porites evermanni [NCBI GCA_942486025.1]
Porites lobata [NCBI GCA_942486035.1]
Acropora digitifera [NCBI GCF_000222465.1]
Acropora millepora [NCBI GCF_013753865.1]
Acropora millepora [Ensemble Amil_v2.1]
Stylophora pistillata [UP000225706]
Stylophora pistillata [NCBI GCF_002571385.2]
Stylophora pistillata [Ensemble Stylophora_pistillata_v1]
Pocillopora verrucosa [NCBI GCF_030620025.1]
Pocillopora meandrina [NCBI GCA_942486045.1]
Pocillopora damicornis [UP000275408]
Pocillopora damicornis [NCBI GCF_003704095.1]
Pocillopora damicornis [Ensembl ASM370409v1]
Actinostola sp. cb2023 [NCBI GCA_033675265.1]
Exaiptasia diaphana [UP000887567]
Exaiptasia diaphana [NCBI GCF_001417965.1]
Exaiptasia diaphana [Ensemble Aiptasia_genome_1.1]
Nematostella vectensis [Ensemble ASM20922v1]
Nematostella vectensis [UP000001593]
Nematostella vectensis [NCBI GCF_932526225.1]
Actinia equina [Ensemble equina_smartden.arro4.noredun]
Actinia tenebrosa [UP000515163]

Actinia tenebrosa [NCBI GCF_009602425.1]
Actinia tenebrosa [Ensembl ASM960242v1]
